# Mitigation of Fetal Irradiation Injury from Mid-Gestation Total Body Radiation with Mitochondrial-Targeted GS-Nitroxide JP4-039

**DOI:** 10.1101/2024.02.13.580105

**Authors:** Yijen L. Wu, Anthony G. Christodoulou, Jan H. Beumer, Lora H. Rigatti, Renee Fisher, Mark Ross, Simon Watkins, Devin R. E. Cortes, Cody Ruck, Shanim Manzoor, Samuel K. Wyman, Margaret C. Stapleton, Eric Goetzman, Sivakama Bharathi, Peter Wipf, Tuantuan Tan, Julie L. Eiseman, Susan M. Christner, Jianxia Guo, Cecilia W. Y. Lo, Michael W. Epperly, Joel S. Greenberger

**Affiliations:** Department of Pediatrics, School of Medicine, University of Pittsburgh, Pittsburgh, PA 15201; Rangos Research Center Animal Imaging Core, Children’s Hospital of Pittsburgh of UPMC, Pittsburgh, PA 15224; Department of Radiological Sciences, David Geffen School of Medicine at UCLA, Los Angeles, CA 90095; Department of Pharmaceutical Sciences, School of Pharmacy, University of Pittsburgh, Pittsburgh, PA 15261; Cancer Therapeutics Program, UPMC Hillman Cancer Center, Pittsburgh, PA 15232; Division of Laboratory Animal Resources (DLAR), University of Pittsburgh, Pittsburgh, PA 15213; Department of Radiation Oncology, School of Medicine, UPMC Hillman Cancer Center, Pittsburgh, PA 15232; Department of Cell Biology, School of Medicine, University of Pittsburgh, Pittsburgh, PA 15260; Department of Biomedical Engineering, Swanson School of Engineering, University of Pittsburgh, Pittsburgh, PA 15261; Department of Chemistry, Kenneth P. Dietrich School of Arts & Sciences, University of Pittsburgh, Pittsburgh, PA 15260; Department of Pharmacology & Chemical Biology, School of Medicine, University of Pittsburgh, Pittsburgh, PA 15213

**Keywords:** total body irradiation, radiation mitigation, fetal, GS-nitroxide, mitochondria, MRI

## Abstract

Victims of a radiation terrorist event will include pregnant women and unborn fetuses. Mitochondrial dysfunction and oxidative stress are key pathogenic factors of fetal irradiation injury. The goal of this preclinical study is to investigate the efficacy of mitigating fetal irradiation injury by maternal administration of the mitochondrial-targeted gramicidin S (GS)- nitroxide radiation mitigator, JP4-039. Pregnant female C57BL/6NTac mice received 3 Gy total body ionizing irradiation (TBI) at mid-gestation embryonic day 13.5 (E13.5). Using novel time- and-motion-resolved 4D *in utero* magnetic resonance imaging (4D-uMRI), we found TBI caused extensive injury to the fetal brain that included cerebral hemorrhage, loss of cerebral tissue, and hydrocephalus with excessive accumulation of cerebrospinal fluid (CSF). Histopathology of the fetal mouse brain showed broken cerebral vessels and elevated apoptosis. Further use of novel 4D Oxy-wavelet MRI capable of probing *in vivo* mitochondrial function in intact brain revealed significant reduction of mitochondrial function in the fetal brain after 3Gy TBI. This was validated by *ex vivo* Oroboros mitochondrial respirometry. Maternal administration JP4-039 one day after TBI (E14.5), which can pass through the placental barrier, significantly reduced fetal brain radiation injury and improved fetal brain mitochondrial respiration. This also preserved cerebral brain tissue integrity and reduced cerebral hemorrhage and cell death. As JP4-039 administration did not change litter sizes or fetus viability, together these findings indicate JP4-039 can be deployed as a safe and effective mitigator of fetal radiation injury from mid-gestational in utero ionizing radiation exposure.

**One Sentence Summary:** Mitochondrial-targeted gramicidin S (GS)-nitroxide JP4-039 is safe and effective radiation mitigator for mid-gestational fetal irradiation injury.

## INTRODUCTION

Radiation mitigators for victims of a radiation terrorist event must be efficacious and safe in all patient populations, including pregnant women and unborn fetuses. Total body irradiation (TBI) can be both lethal and teratogenic for unborn fetuses, in particular, the developing nervous system. Integration of the developing fetus and the maternal environment (*1–4*) greatly impacts fetal development. The adverse effects of TBI on developing fetuses depend on the stages of embryogenesis and fetal development (*5–8*). In both animal models and humans, ionizing radiation induces significant developmental abnormalities during mid-gestation (*9, 10*). Mid-gestational radiation toxicity has been shown to involve the central nervous system in both non-human primates (*11, 12*) and humans (*13–16*). In particular, the developing hippocampus (*17–20*) and neonatal vasculature (*11, 12, 21*) have been shown to be sensitive targets for radiation damage (*5, 22*). In addition, therapeutic radiotherapy for cancers has been well-documented to cause radiation damage to neurocognitive functions in both the pediatric and adult human brain (*23–26*).

TBI is known to cause oxidative stress (*27–30*) with increased release of reactive oxygen species (ROS). TBI also can lead to mitochondrial dysfunction, including changes in the mitochondrial genome, as well as activation of mitophagy and degenerative processes (*31–35*). Therefore, mitochondria-targeting ROS scavengers are promising agents to mitigate irradiation injury (*36*). We have previously shown that the mitochondria-targeting gramicidin S (GS)- nitroxide, JP4-039 (*37*), can augment the potentially lethal damage repair (PLDR) in cells after irradiation (*38*) and in mice after TBI (*39–43*). However, it is not known if JP4-039 can mitigate irradiation injury in developing fetal brains after TBI in pregnant females.

We investigated the effects of 3.0 Gy TBI on embryonic day E13.5 mid-gestation mouse fetuses, and the impact of maternal administration of JP4-039 one day after irradiation, on E14.5. To evaluate the efficacy of JP4-039, we first measured the placental-fetal drug transit (*44*) to show that JP4-039 can pass through the placental barrier to reach fetuses. Leveraging a novel time-and-motion-resolved 4D *in utero* magnetic resonance imaging (4D-uMRI) method, we evaluated fetal brain tissue integrity, cerebral hemorrhage, and cerebrospinal fluid (CSF). The *in vivo* 4D-uMRI findings were validated by *ex vivo* high-resolution MRI, histopathology, and immunofluorescence microscopy. Furthermore, we used the novel 4D Oxy-wavelet MRI to probe *in vivo* mitochondrial function in fetal brains, which was validated with *ex vivo* Oroboros mitochondrial respirometry. The results showed robust mitigation of irradiation-induced fetal brain injury in mid-gestation by administration of JP4-039 to pregnant female mice, and no adverse effects of the treatment.

## RESULTS

### Fetal Total Body Irradiation in Mid-Gestation in Mice

Previous studies (*45*) in mice demonstrated prenatal lethality and severe teratogenicity with modest total body irradiation (TBI) doses during early gestation (equivalent to human 1^st^ trimester), indicating early conceptuses are highly vulnerable to TBI. The objective of this study is to evaluate amelioration of TBI effects to ultimately improve fetal survival with mid-gestation (equivalent to human later 1^st^ trimester and the beginning of the 2^nd^ trimester) TBI in mice on embryonic day (E13.5). At this development stage, the 4-chamber heart is formed, and thus would allow the assessment for impaired cardiovascular function and also possible presence of structural heart defects that can cause prenatal lethality.

We previously measured irradiation dose responses of 1.0, 3.0, 5.0, and 7.0 Gy TBI on pregnant female mice on E13.5 and found a viability index (percentage of animals that survive 4 days or longer) of 95%, 31%, 0%, and 0%, respectively, with 100% lethality at doses above 3.0 Gy (*44*). To test the efficacy of JP4-039 in ameliorating these effects in the current study, a 3.0 Gy irradiation was chosen. This is predicted to yield liveborn pups, but at a significantly lower survival rate than in sham controls.

With 3 Gy TBI on E13.5, the gestation index (percentage of pregnancies resulting in live litters), viability index (percentage of animals that survive 4 days or longer), and lactation index (percentage of animals alive at 4 days that survived the 21-day lactation period) are 100%, 31%, and 7%, respectively (*44*). To investigate the possible causes of neonatal lethality, we further used in utero ultrasound imaging to assess fetal heart structure and function with interrogation of fetuses at E15.5. This analysis showed no structural heart defects nor changes in fetal cardiovascular functions (Supplemental Figure). Therefore, the observed neonatal lethality was not due to cardiovascular defects.

### JP4-039 can effectively cross placental-fetal barrier to reach fetuses

As a potential agent for ameliorating fetal irradiation injury via maternal administration, JP4-039 needs to be able to cross the placental-fetal barrier. Hence, the presence of JP4-039 in the placenta and fetuses was evaluated on E14.5 in pregnant C57BL/6NTac mice using validated liquid chromatography-tandem mass spectroscopy (LC-MS) (*46*). One day after 3.0 Gy TBI on E13.5, JP4-039 was administered to pregnant female mice on E14.5. Maternal and fetal tissues were harvested 30 minutes after JP4-039 administration. The concentrations of JP4-039 in maternal plasma (Fig.1A), maternal brains (Fig.1B), and placentas (Fig.1C), as well as fetal brains (Fig.1D) and fetal bodies (Fig.1E) were quantified in 3 litters per group. The number of fetuses per litter varied from 5 to 10 per pregnant female mouse, with 0 to 4 resorptions. The litter sizes did not appear to be different between TBI (7.00±0.58) and sham litters (6.67±0.88). Thirty minutes after dosing, JP4-039 was found in fetal brains (Fig.1D), fetal bodies (Fig.1E), and placenta (Fig.1C), indicating JP4-039 can successfully cross the placental-fetal barrier and reach the fetus. The concentration of JP4-039 in fetal brain (Fig.1D), the fetal body (Fig.1E), and placenta (Fig.1C) was not different among litters with or without irradiation. This suggests that irradiation injury from 3 Gy TBI did not abolish JP4-039 transport to fetuses. Although the JP4-039 concentration in maternal plasma (Fig.1A) appeared to be lower in irradiated (IR) mice, it did not impact the amount of JP4-039 being transported across the placenta to reach the fetus (Fig.1D, E). Most importantly, the JP4-039 concentration in fetal brain (Fig.1D) was similar to that of fetal body (Fig.1E) after irradiation. Furthermore, the JP4-039 concentration in both fetal brain (Fig.1D) and fetal body (Fig.1E) after irradiation were still higher than that in maternal brains (Fig.1B). Our data show that JP4-039 can be effectively transported across the placental-fetal barrier to reach the fetus within 30-minutes post-dosing, and this transport mechanism is not altered by 3 Gy TBI. Therefore, JP4-039 can potentially be an effective amelioration agent for fetal irradiation injury via maternal administration.

**Figure 1.**
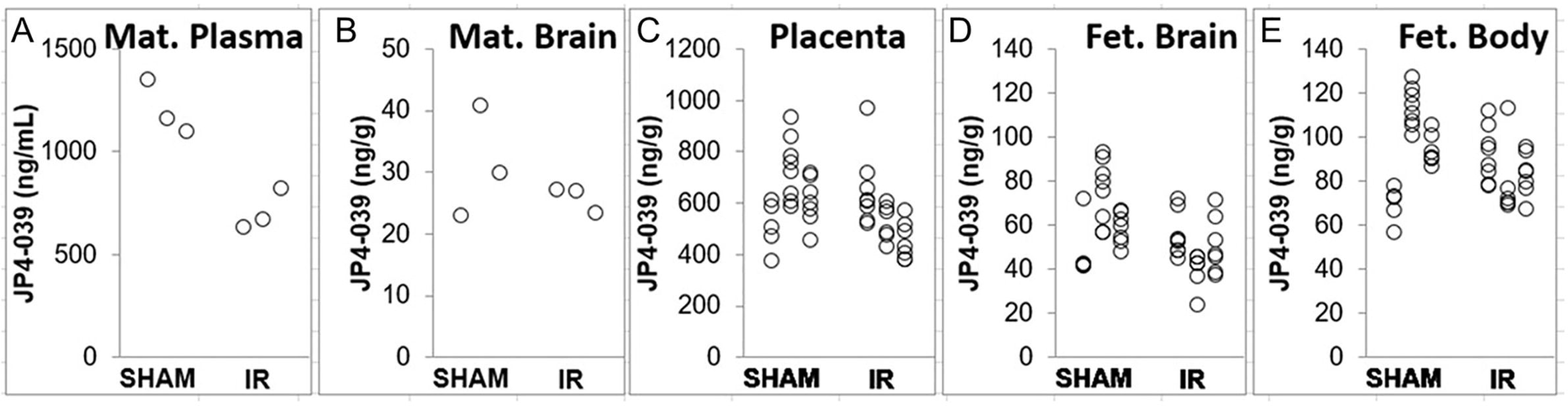
Quantitation of Maternally Administered JP4-039 in Placentae and Fetal Brains on E14.5 with LC-MS. JP4-039 was quantified with LC-MS on E14.5 at 30 minutes after maternal administration. IR – with 3 Gy irradiation. Sham – with 0 Gy irradiation on E13.5. (A) maternal mice in the sham group had significantly higher JP4-039 plasma concentrations (mean ± SD: 1203.3 ± 130.5) than the IR group (mean ± SD: 706.7 ± 99.6; p=0.0063). (B) maternal mice in the sham group did not have significant different JP4-039 brain exposure (mean ± SD: 31.3 ± 9.0) than the IR group (mean ± SD: 25.8 ± 2.2; p=0.3629). (C) pups in sham group did not have significantly different JP4-039 placenta exposure (mean ± SD: 633.2 ± 136.9) than the IR group (mean ± SD: 547.9 ± 132.5; p=0.3387). (D) pups in sham group did not have significant different JP4-039 brain exposure (mean ± SD: 62.3 ± 15.9) than the IR group (mean ± SD: 49.3 ± 12.0; p=0.1734). (E) pups in sham group did not have significant different JP4-039 exposure in the rest tissues (mean ± SD: 95.4 ± 19.2) than the IR group (mean ± SD: 85.1 ± 13.8; p=0.4984).

### Improved Survival of JP4-039 Treated Fetal Mice after Mid-gestation TBI

We examined the effect of JP4-039 on litter sizes and survival of mid-gestational fetal mice (Table 1). Litter size at birth from the unirradiated sham controls (0 Gy) was 8.4/litter with a 98 ± 5 % survival to weaning. Litter size from the unirradiated vehicle control females (0 Gy + JP4-039) that received JP4-039 was 7.3 with an 85 ± 17% survival to weaning, which is statistically not significantly different from those of the sham unirradiated control females. Pregnant females receiving 3.0 Gy on E13.5 had a litter size of 7.7/litter, which is also not different from the unirradiated sham (0 Gy) or vehicle (0 Gy + JP4-039) controls. However, while most pups irradiated on E13.5 were born alive they did not survive past day one. There were only 5 out of 71 (7 ± 19%) pups that survived till weaning, and less than 1% survived till 3 months. In contrast, pregnant females irradiated on E13.5, then received JP4-039 treatment on E14.5 (3 Gy + JP4-039), had similar litter sizes of 7.8/litter at birth, their pubs showed significantly improved survival with 44 out of 78 (56 ± 40%) surviving till weaning (*p* = 0.0027) (Table 1). Most JP4-039 treated survivors lived to adulthood. Our data demonstrate that a single maternal administration of JP4-039 after mid-gestation TBI dramatically improved postnatal survival.

**Table 1.**
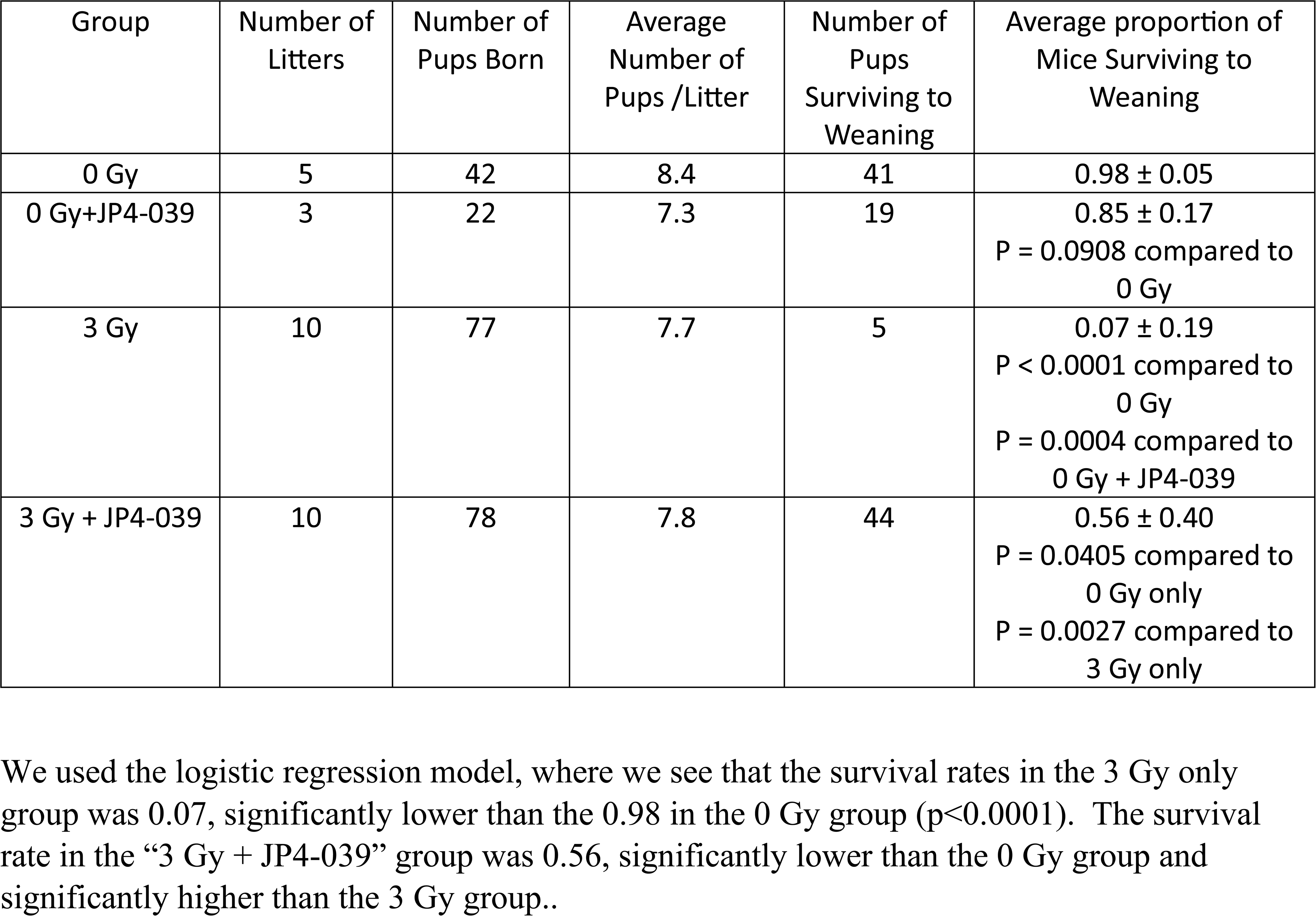

### JP4-039 Reduced TBI-induced Fetal Brain Tissue Loss and Cerebral Hemorrhage

To examine the impact of JP4-039 on potential rescue of TBI-induced fetal brain injury, we leveraged a novel tome-and-motion-resolved 4D *in utero* MRI (4D-uMRI) capable of simultaneous *in utero* imaging of multiple mouse fetuses with high spatiotemporal resolutions without motion artifacts. 4D-uMRI on E16.5, which was three days after the 3 Gy TBI, showed the irradiated fetuses (Fig.2B) had extensive cerebral hemorrhage, loss of brain tissues, and enlarged ventricles with excessive accumulation of cerebrospinal fluid (CSF). In contrast, the irradiated litters that received maternal administration of JP4-039 on E14.5 (Fig.2C) showed lesser degrees of brain tissue damage with less cerebral hemorrhage and less CSF volume increase, and improved brain tissue integrity. Fetal brain tissue volumes (Fig.2 D, G), CSF volumes (Fig. 2E, H), and cerebral hemorrhage volumes (Fig.2 F, I) were quantified for each group. The irradiated fetuses were noticeably smaller, and thus the volumes were quantified in both ways, either by direct measurement (Fig.2 D-F) or normalized with respect to total brain volume (Fig.2 G-I). Volumetric analysis of the *in vivo* 4D-uMRI found that fetuses treated with 3 Gy TBI on E13.5 (Fig.2 D-I, red, 3 Gy, n = 29) had significant reduction in brain tissue volume (Fig.2 D, G), higher CSF volume (Fig. 2E, H), and more cerebral hemorrhage (Fig.2 F, I) as compared to that of sham controls (Fig.2 D-I, blue, 0 Gy, n = 25). In contrast, the irradiated fetuses that received maternal administration of JP4-039 on E14.5 (Fig.2 D-I, green, 3 Gy + JP4, n = 26) had increased brain tissue volume (Fig.2 D, G), lower CSF volume (Fig. 2E, H), and less cerebral hemorrhage (Fig.2 F, I). Maternal administration of JP4-039 greatly reduced TBI-induced cerebral hemorrhage, hydrocephalus, and brain tissue loss.

**Figure 2.**
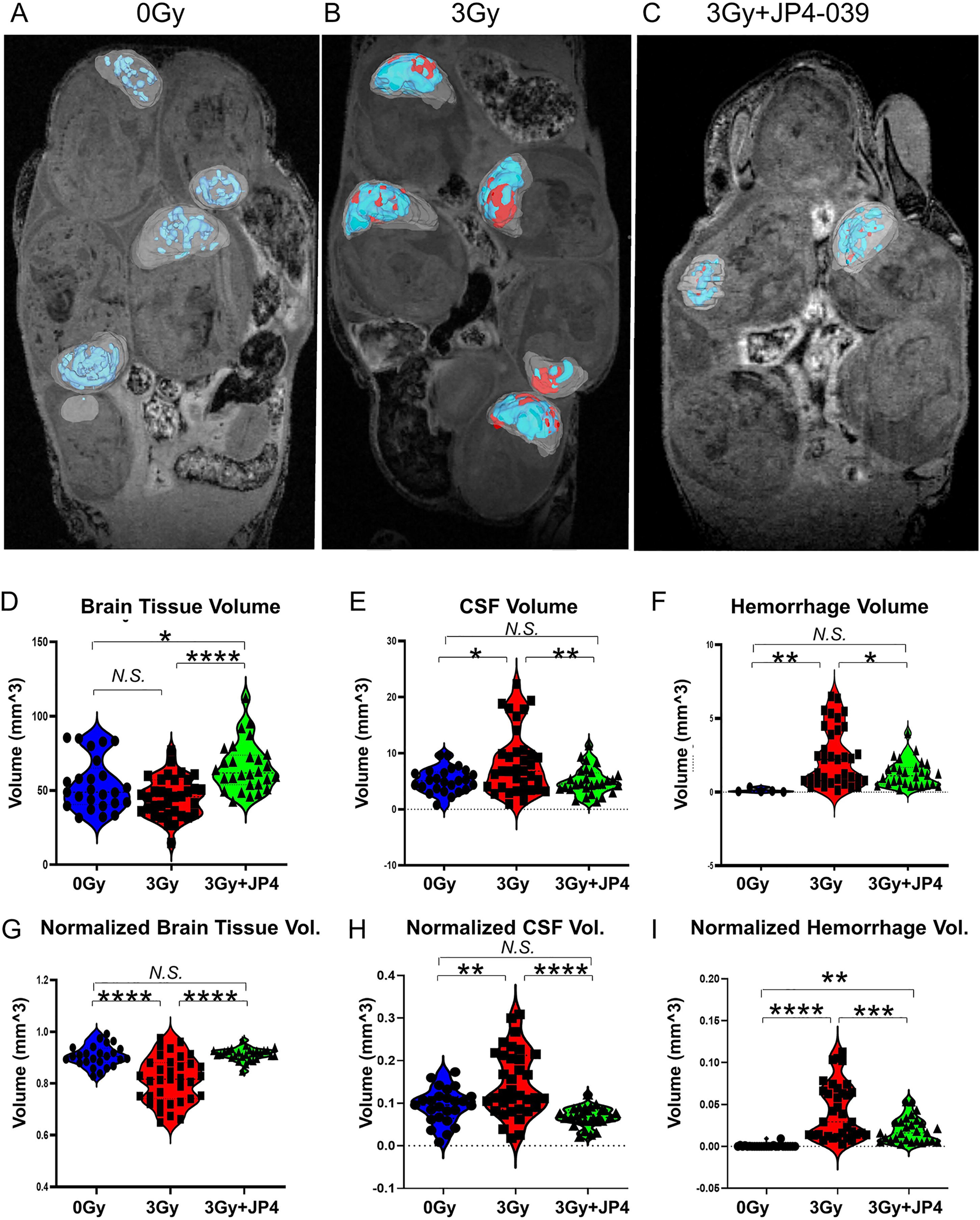
Maternal administration of JP4-039 on E14.5 after 3Gy TBI on E13.5 could improve cerebral tissue preservation, reduce cerebral hemorrhage, and excessive cerebrospinal fluid (CSF) from 3Gy irradiation, evaluated by *in vivo* 4D-uMRI on E16.5. (A-C) *In vivo* 4D-uMRI on E16.5 of a naïve control litter (A, 0Gy), a litter received 3Gy TBI on E13.5 (B), and a litter received 3Gy on E13.5 and JP4-039 on E14.5 (C). One plane of the 3D stack is shown for each litter. Segmentation masks: gray – fetal brain; blue - cerebrospinal fluid (CSF), and red - cerebral hemorrhage. (D-I) Volumetric analysis of fetal brain tissue volume (D), CSF volume (E), cerebral hemorrhage volume (F), normalized fetal brain tissue volume (G), normalized CSF volume (H), and normalized cerebral hemorrhage volume (I). Blue – naïve control (0Gy, n = 25), red −3Gy irradiated (3Gy, n = 29), and green – 3 Gy with JP4-039 treated (3Gy + JP4, n = 26). *N.S.:* not significant; *: *p* < 0.05; **: *p* < 0.005; ***: *p* < 0.0005; ****: *p* < 0.0001.

### *Ex Vivo* High Resolution MRI Confirmation of *In Vivo* 4D-uMRI Findings

The *in vivo* 4D-uMRI findings were confirmed by additional *ex vivo* high-resolution MRI (Fig.3). This showed extensive cerebral hemorrhage (Fig.3 A-C, pink “H”) along the middle and anterior cerebral arteries, sub-arachidonic space, and around the superior sagittal sinus. The lateral ventricles (Fig.3 A-C, blue “V”) were greatly enlarged, but not the 3^rd^ nor the 4^th^ ventricles. The cortical tissues were greatly atrophied or never developed. The brain stem, olfactory bulb, sub-cortical areas, and spinal cord appeared to be morphologically normal. The gross morphology of the visceral organs, such as the heart, lung, liver, stomach, and kidneys also appeared to be largely unchanged. On the other hand, irradiated fetuses that received JP4-039 on E14.5 (Fig. 3 D-F) showed lesser degree of irradiation damage, with less cerebral hemorrhage (Fig. 3 D-F, pink “H”), smaller lateral ventricles (Fig. 3D-F, blue “V”), and more preserved cortical tissues.

**Figure 3.**
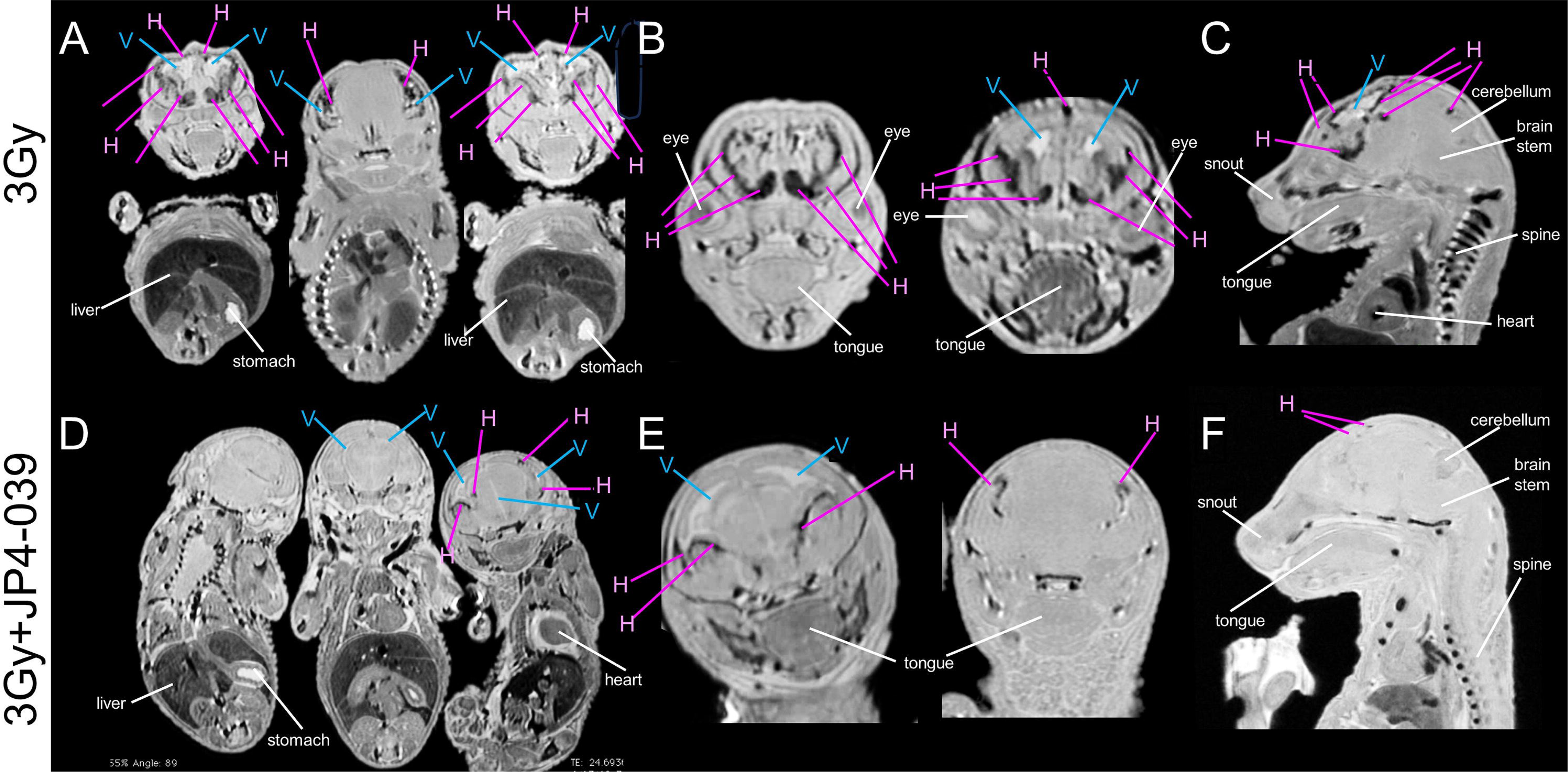
*Ex vivo* high-resolution MRI validating *in vivo* 4D-uMRI evaluation of fetuses harvested on E16.5. (A-C) Fetuses received 3Gy TBI on E13.5, (D-F) Fetuses received 3Gy TBI on E13.5 and JP4-039 on E14.5. (A,D) coronal view; (B, E) axial view; (C, F) sagittal view. Blue lines labeled with “V” point to ventricles, whereas pink lines labeled with “H” point to cerebral hemorrhage.

### *Ex Vivo* Histopathologic Confirmation of MRI Findings

The H&E histopathology (Fig.4) on E18.5 fetal brains confirmed the *in vivo* and *ex vivo* MRI observations. 3 Gy TBI on E13.5 resulted in thinning of cerebral cortex (Fig.4B) and enlarged ventricles on E18.5. In some cases, the lateral ventricles of JP4-039-treated irradiated animals (Fig.4F) are less dilated and comparable to control animals (Fig.4A). Irradiated animals without treatment showed evidence of cerebral vascular damage (Fig.4E, H), including vessel wall thickening, basophilia, and loss of endothelial cells. Blood vessels in the meninges of irradiated animals (Fig.4E, H) are consistently more dilated and congested compared to those of the control (Fig.4D, G). Damaged vessels (Fig.4H, arrow) and extra-vascular red blood cells (RBC) were found in fetal brains with 3 Gy TBI, confirming the *in vivo* and *ex vivo* MRI findings. These vascular changes were not seen in control animals (Fig.4 D, G) and less in irradiated animals treated with JP4-039 (Fig.4 F, I). Cerebellum (Fig.3K) in the 3 Gy TBI treated mice lost the normal cerebellar structure with its cerebellar folia, fissures, and layered organizations seen in the controls (Fig.3J). Preservation of the cerebellar folia/fissures and cell layers were observed in some of the JP4-039 treated animals (Fig.3L).

**Figure 4.**
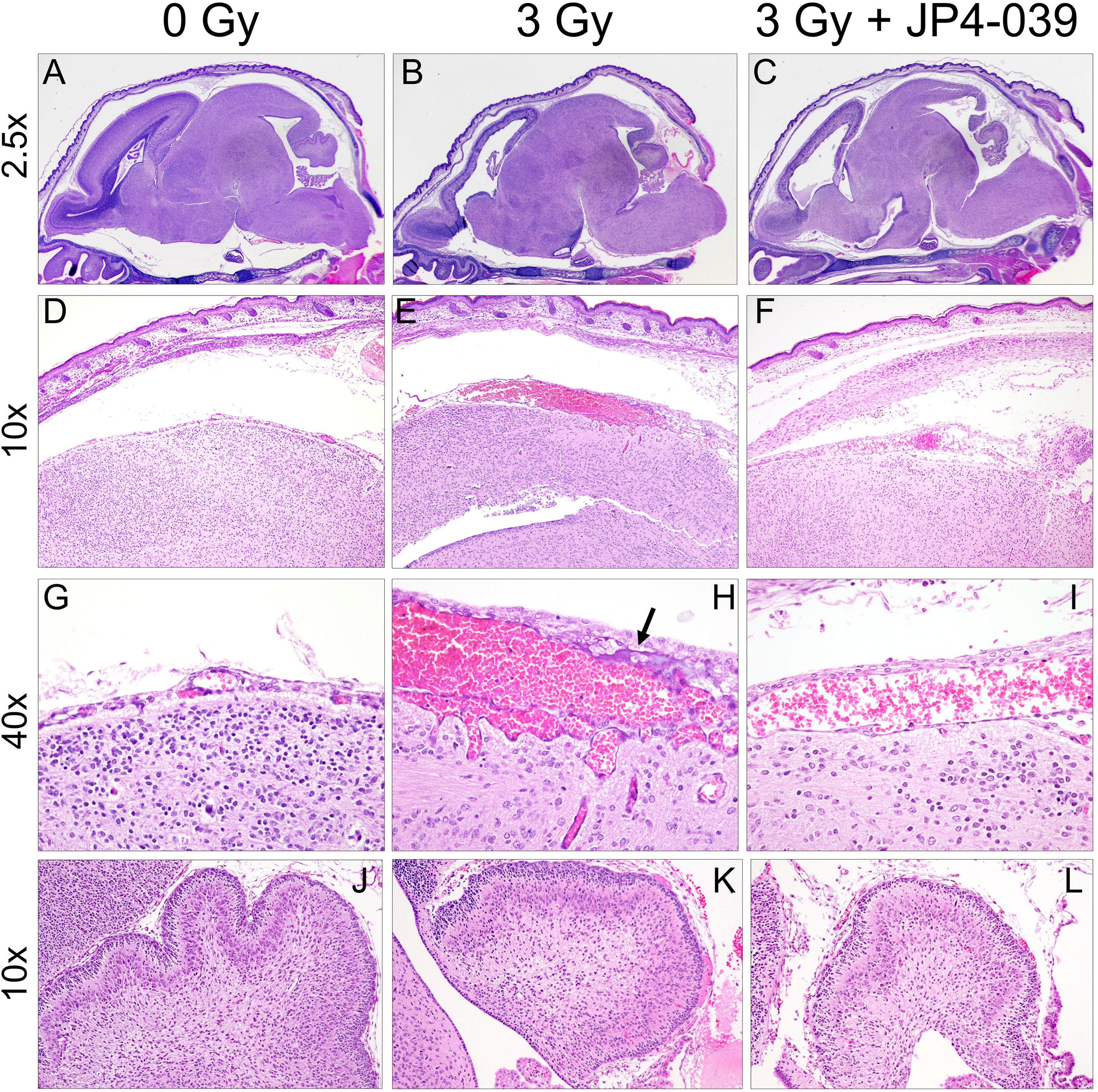
H & E histopathology of fetal brains harvested on E18.5 from (A, D, G, J) a naïve control, (B, E, H, K) a fetus received 3 Gy TBI on E13.5, and (C, F, I, L) a fetus received 3Gy TBI on E13.5 and JP4-039 on E14.5. (D-F) showing 10X magnification. (G-I) showing 40X magnification. Significant congestion is evident in cerebral capillaries of 3 Gy TBI and 3 Gy TBI + JP4-039 brains (E, F, H, L) compared to the naïve control with no congestion (D, G). The arrow points to a necrotic capillary wall present only within the congested region of the 3 Gy TBI brain. (J-L) showing 10X of cerebellum. Loss of cerebellar folia and fissures in 3 Gy TBI and 3Gy TBI + JP4-039 sections is evident (K, L) when compared to naïve control (J).

### JP4-039 Can Reduce TBI-induced Apoptosis in Fetal Brains

Immunofluorescent microscopy with TUNEL staining (Fig.5) showed elevated apoptosis (Fig.5B pink and Fig. 5E) in fetal brains that received 3 Gy on E13.5 but not in controls (Fig.5 AD). The areas with high apoptosis are in the vicinity of cerebral hemorrhage observed with MRI (Fig.3). The irradiated fetal brains that received JP4-039 on E14.5 (Fig. 3 C, F) showed markedly reduced apoptosis relative to irradiated animals. Our data show maternal administration of JP4-039 one day after 3 Gy TBI in the mid-gestation reduced TBI-induced apoptosis.

**Figure 5.**
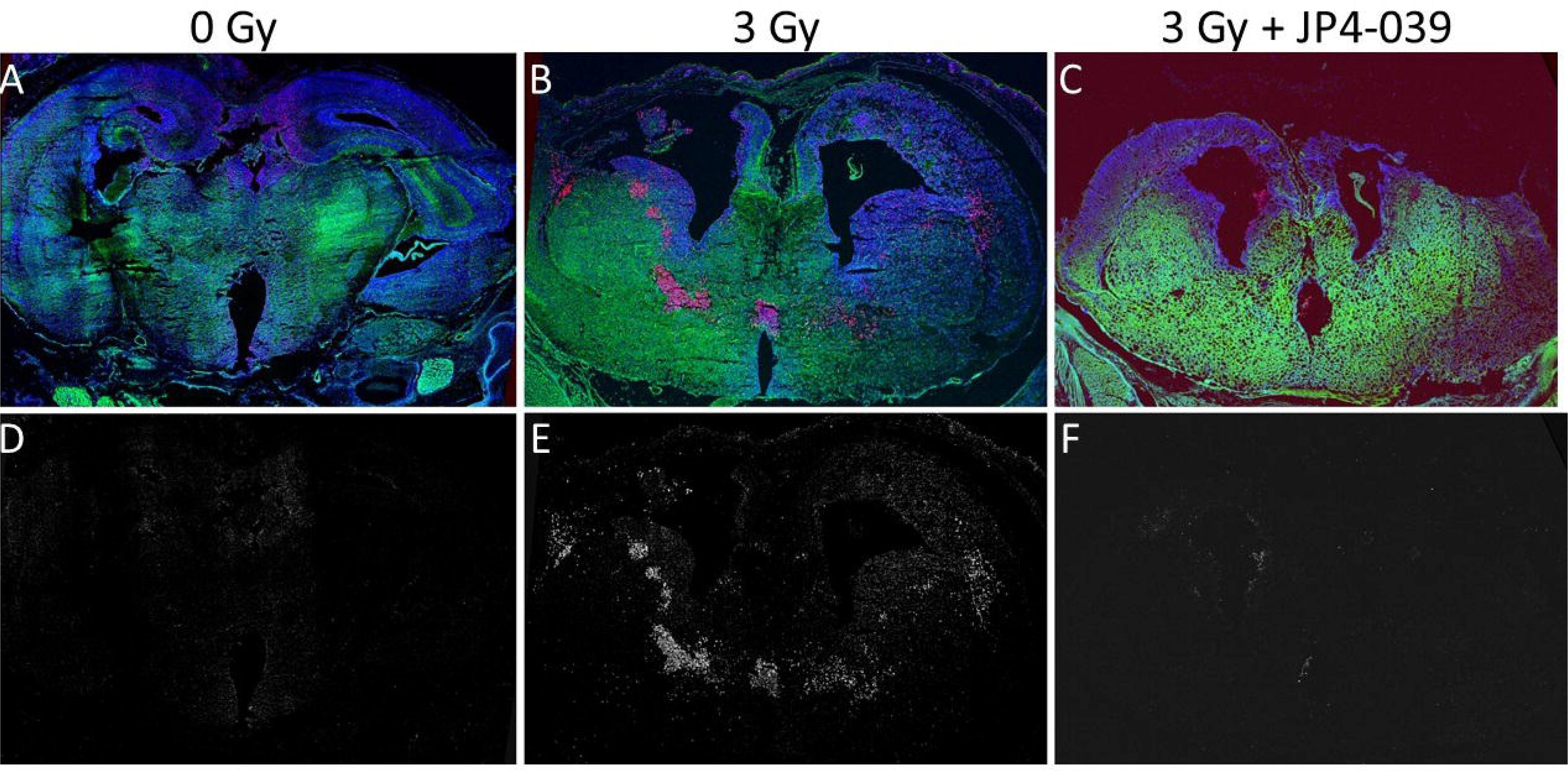
Maternal administration of JP4-039 on E14.5 after 3Gy TBI on E13.5 could reduce the elevated apoptosis from 3Gy irradiation evaluated by fluorescent immunohistochemistry. (A, D) a naïve control, (B, E) a fetus received 3Gy TBI on E13.5, and (C, F) a fetus received 3Gy TBI on E13.5 and JP4-039 on E14.5. (A-C) composite of immunofluorescent microscopy. Pink – TUNEL staining for apoptosis. Blue – DAPI for nuclei. (D-F) single channel fluorescence of TUNEL staining for apoptosis.

### JP4-039 Can Reduce TBI-induced Mitochondrial Dysfunction in Fetal Brains

Acute (*47–51*) adaptation to hypoxia requires mitochondria-controlled signaling (*52, 53*) for transient, reversible, compensatory activation of the respiratory chain to maintain oxygen homeostasis, a major mechanism of immediate adaptation to hypoxia. We leveraged this property to probe mitochondrial function *in vivo* using 4D Oxy-wavelet MRI with oscillating short bursts (3 min) of hypoxia (10% O_2_) challenges (Fig.6A). Using this novel method, we successfully probed fetal brain mitochondrial function *in utero* after 3 Gy TBI with maternal inhalation of an oscillating hypoxia challenge (Fig.6A). Upon oscillating hypoxia challenges, normal fetal mouse brains with intact mitochondrial functions can overcome the bursts of hypoxia (Fig.6B, orange periods) to quickly adjust to a higher oxygenation state after the initial transient functional MRI (fMRI) blood-oxygen-level-dependent (BOLD) signal drop, indicating intact *“brain sparing”* capability, with most voxels showing higher than baseline BOLD levels (Fig.6E, green). The summation of all the BOLD voxels during the hypoxia period, reflecting the overall fetal brain oxygenation, was positive, higher than the baseline (Fig.6H, green). Conversely, fetal brain subjected to a 3 Gy irradiation with compromised mitochondrial function showed a persistently decreased BOLD signal (Fig.6C), indicating an inability to cope with hypoxia, with most voxels showing BOLD levels lower than baseline (Fig.6F, red). The summation of all the BOLD voxels during the hypoxia period, reflecting the fetal brain oxygenation, was negative, lower than the baseline (Fig.6I, red). In contrast, some of the irradiated fetuses that received maternal administration of JP4-039 (Fig.6D) were able to respond similarly to normal ones, quickly adjusting to a higher oxygenation state after the initial transient BOLD signal drop, and with most voxels showing higher than baseline BOLD levels (Fig.6G, green). The summation of all the BOLD voxels during hypoxia period, reflecting the fetal brain oxygenation, was positive, higher than the baseline (Fig.6J, green). This is indicative that JP4-039-treated fetal brain had a high mitochondrial function. Evaluation of the BOLD summation, reflecting total fetal brain oxygenation status during the hypoxia challenge, across 5 litters (Fig.6K) showed irradiated fetal brain (Fig.6K, red, 3 Gy, n = 29) had lower BOLD summation compared to that of the control brain (Fig.6K, blue, 0 Gy, n = 25). This indicates that irradiated fetal brain is unable to maintain oxygen homeostasis during hypoxia challenges, suggesting compromised mitochondrial function. JP4-039 treatment rescued some fetuses from this irradiation-induced brain mitochondrial injury. Some JP4-039 treated fetuses (Fig.6K, green, 3 Gy + JP4, n = 26) showed high brain BOLD summation, reflecting total oxygenation in fetal brain during the hypoxia challenge, suggesting elevated mitochondrial function, even as some treated fetuses continued to show low brain mitochondrial function. Overall, combining all JP4-039 treated fetuses (Fig.6K green) still showed higher functions compared to the untreated, irradiated counterparts (Fig.6K red).

**Figure 6.**
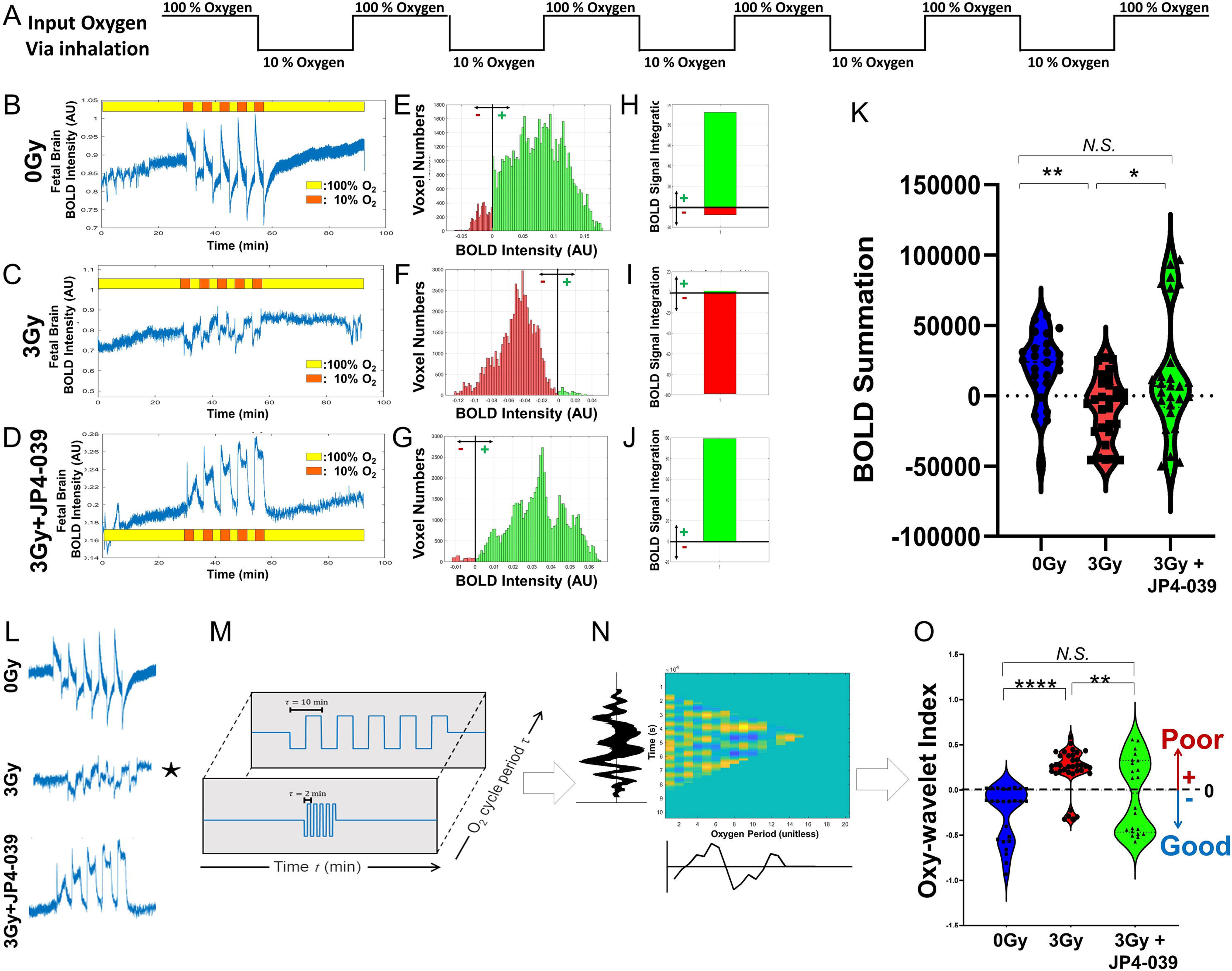
*In vivo* 4D Oxy-wavelet MRI probing mitochondrial dysfunction in fetal irradiation injury on E16.5. (A) Input oxygen wave via maternal inhalation. Oxygen supply for the pregnant female via a nose cone is alternating between 100% and 10% oxygen every 3 minutes. (B-D) Temporal kinetics of BOLD signals in fetal brains. Yellow: 100% oxygen. Orange: 10% oxygen. (E-G) Histogram pixels during hypoxia periods. Green-time pixels of BOLD signal higher than baseline; Red-time pixels of BOLD signal lower than baseline. (H-J) Summation of BOLD histogram areas. Green-time pixels of BOLD signal higher than baseline; Red-time pixels of BOLD signal lower than baseline. (B, E, H) from a naïve control fetal brain. (C, F, I) from a fetal brain that received 3Gy TBI on E13.5. (D, G, J) from a fetal brain that received 3Gy TBI on E13.5 and JP4-039 on E14.5. (K) Composite BOLD summation during the hypoxia periods for blue – naïve control (0Gy, n = 25), red −3Gy irradiated (3Gy, n = 29), and green – 3 Gy with JP4-039 treated (3Gy + JP4, n = 26). *N.S.:* not significant; *: *p* < 0.05; **: *p* < 0.005. (L-O) Computerized waveform analysis for 4D Oxy-wavelet MRI. (L) Example region-averaged BOLD response waveforms for each group: naïve (top), irradiation exposed (middle), irradiation exposed and treated with JP4-039 (bottom). (M) Iterative correlation with the oxygen event waveform at various oxygen flipping periods, Tau. This allows for automatic detection of the appropriate oxygen event scheme via maximization of the cross-correlation. (N) Example time and oxy-wavelet space, with the oxygen marginal and time marginal displayed across the x- and y-axis. (O) Violin plot comparing the final Oxy-wavelet index across the three animal groups: Blue – naïve control (0Gy, n = 25), red −3Gy irradiated (3Gy, n = 29), and green – 3 Gy with JP4-039 treated (3Gy + JP4, n = 26). *N.S.:* not significant; *: *p* < 0.05; **: *p* < 0.005; ****: *p* < 0.0001. Negative Oxy-wavelet index indicates good mitochondrial functions enabling active compensation during hypoxia challenges, whereas positive Oxy-wavelet index indicates poor mitochondrial functions, passive inability to compensate for hypoxia challenge.

We further used the Oxy-wavelet index to track mitochondrial function (Fig.6O) in the normal and irradiated mice, with and without JP4 treatment. A positive index indicated poor mitochondrial function that cannot compensate for the hypoxia challenge to maintain oxygen homeostasis, while negative Oxy-wavelet index indicates good mitochondria function capable of active compensation of the hypoxia challenge to maintain oxygen homeostasis (see Methods). As expected, the Oxy-wavelet waveform analysis (Fig.6O) showed the sham controls (Fig. 6O, blue, 0Gy, n = 25) had a negative Oxy-wavelet index, indicating intact mitochondrial function, capable of actively maintaining oxygen homeostasis during the hypoxia challenge. The irradiated fetal brains (Fig. 6O, red, 3Gy, n = 29) showed mostly positive Oxy-wavelet index, indicating poor mitochondrial functions unable to maintain oxygen homeostasis during the hypoxia challenge. In contrast, the fetal brains exposed to irradiation and treated with JP4-039 (Fig. 6O, green, 3Gy + JP4, n = 26) had a binodal Oxy-wavelet response that were on average statistically significantly better than the irradiated group, showing no difference compared to the control group. The *in vivo* 4D Oxy-wavelet MRI showed TBI on E13.5 greatly compromised the fetal brain mitochondrial respiratory function, but maternal administration of JP4-039 on E14.5 improved the mitochondrial respiratory function in most, although not all fetal brains.

### Biochemical Validation of Mitochondrial Function Assessment by 4D Oxy-wavelet MRI

*In vivo* 4D Oxy-wavelet MRI evaluation of fetal brain mitochondrial function was validated with *ex vivo* Oroboros mitochondrial respirometry (Fig.7). Immediately following the 4D Oxy-wavelet MRI on E16.5, mitochondria were isolated from the fetal brain lysates. Mitochondrial respiratory chain function was evaluated by the step-wise addition of substrates. All three groups showed similar baseline respiratory activities (Fig.7B, G, L). The irradiated group (Fig.7A blue and Fig.7B-E, red) showed reduced respiratory chain complex I and pyruvate-dependent activities. Complex II activity was not significantly different. With treatment of JP4-039, some of the fetal brains showed improved functions while others did not show improvements with 4D Oxy-wavelet MRI (Fig.6O green). Fast *in vivo* 2D MRI was used to parcellate JP4-039 treated fetal brains into high and low injury categories. The JP4-039 treated fetal brains that showed improvement with the *in vivo* MRI also showed improvement with the *ex vivo* Oroboros analysis (Fig.7F red and Fig.7 G-J green), mostly on the complex I and pyruvate-related activities. On the other hand, fetal mice treated with JP4-039 that did not show improvement with *in vivo* MRI also showed no improvement in the *ex vivo* Oroboros analysis (Fig.7K red and Fig.7 L-O, green). The *ex vivo* Oroboros mitochondrial respirometry confirmed the *in vivo* 4D Oxy-wavelet MRI findings that JP4-039 could rescue mitochondrial dysfunctions in irradiated fetal brains. The fetal brains with higher mitochondrial respiratory functions had lower fetal brain injury.

**Figure 7.**
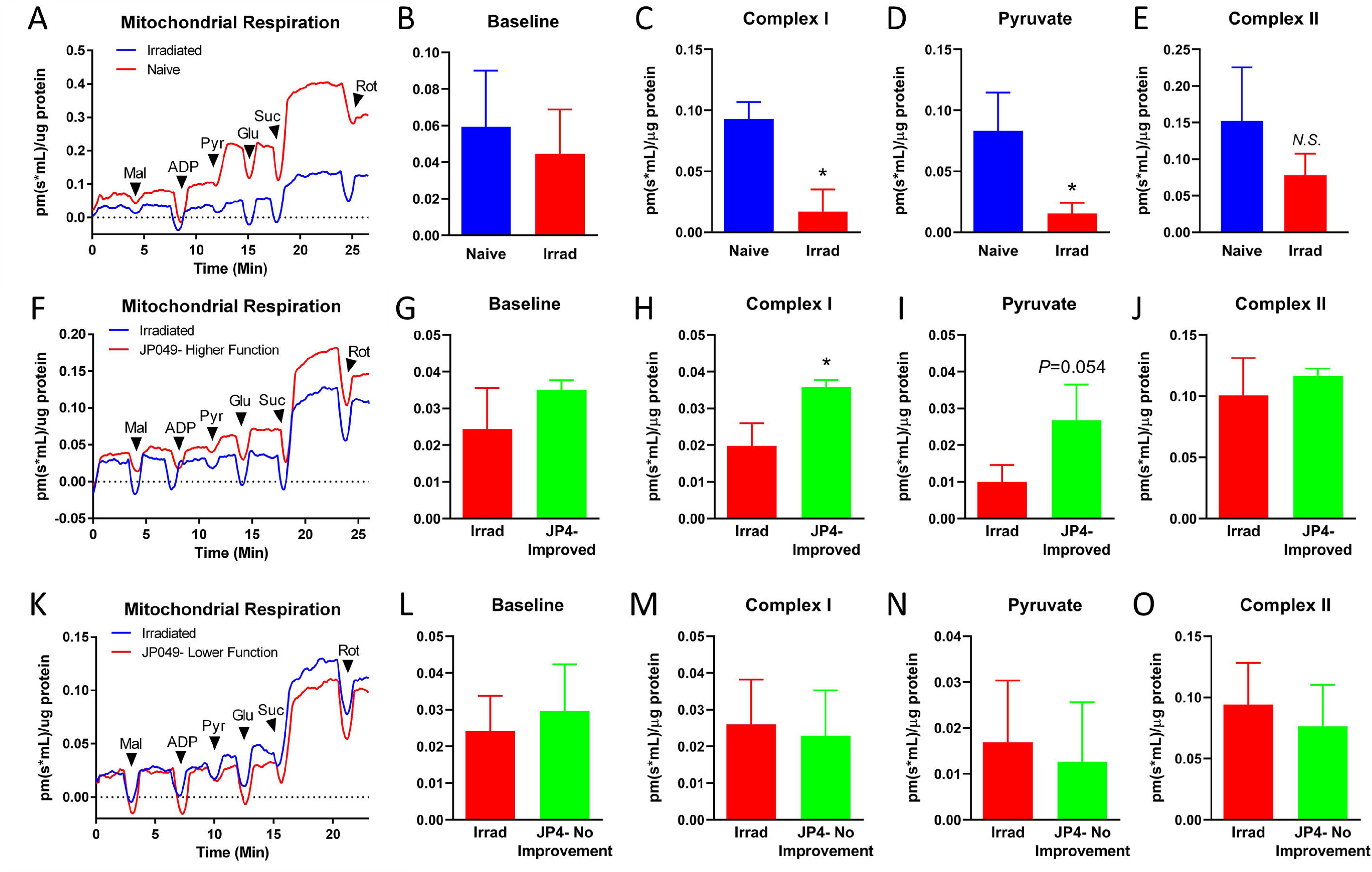
Mitochondrial targeting JP4-039 attenuation of irradiation-caused mitochondrial dysfunction, evaluated by *ex vivo* Oroboros high-resolution respirometry of isolated mitochondria from fetal brains on E16.5. *Oroboros* mitochondrial respirometry were conducted with isolated mitochondria from fetal brain lysates harvested on E16.5. Substrates and inhibitors were added stepwise: cytochrome-C (CytC), malate (Mal), Pyruvate (Pyr), glutamate (Glu), succinate (Suc), rotenone (Rot). (A-E) mitochondrial respiration compared between irradiated and naïve. (F-J) mitochondrial respiration compared between irradiated and JP4-039 treated that showed high improved function with 4D Oxy-wavelet MRI. (K-O) mitochondrial respiration compared between irradiated and JP4-039 treated that showed low function with 4D Oxy-wavelet MRI. (A) blue-irradiated, red-naïve; (F) blue-irradiated, red-irradiated with JP4-039, showing improved high function with 4D Oxy-wavelet MRI; (K) blue-irradiated, red-irradiated with JP4-039, but showing low function with 4D Oxy-wavelet MRI. (B-E, G-J, L-O) group averaged (n=5) comparison. Blue: naïve control, red: irradiated, that received 3Gy TBI on E13.5. Green – JP4-treated, that received 3Gy TBI on E13.5 and JP4-039 on E14.5. (G-J) JP4-039 treated that showed improved function with 4D Oxy-wavelet MRI. (L-O) JP4-039 treated that showed low function with 4D Oxy-wavelet MRI. (B, G, L) baseline respiration (no exogenous substrates), (C, H, M) mitochondrial respiratory chain Complex I respiration (State 3 respiration on malate, glutamate, and pyruvate), (D, I, N) the individual response to pyruvate, and (E, J, O) mitochondrial respiratory chain Complex II respiration (State 3 respiration on succinate). *: *p* < 0.05.

## DISCUSSION

Our study showed 3 Gy TBI on E13.5 fetal mice did not cause any detectable damage to the heart or placenta. At this developmental stage, the 4-chamber heart and fetoplacental circulation have already been established. Neither fetal heart structure nor function was impacted, and the irradiated placenta was still capable of transporting JP4-039 to fetuses, including to the fetal brain. In contrast to the heart, severe fetal brain injury was observed that included cerebral hemorrhage and excessive CSF accumulation with hydrocephalus observed without obstruction or narrowing of the cerebral aqueduct, ie. nonobstructive hydrocephalus. Brain development was disrupted with defects in cortical development. As cortical neurogenesis (*54*), neuronal migration, and maturation in the mouse brain are initiated around E12.5, E14.5, and E16.5, respectively, and continuing after birth, perturbation of brain development and brain damage may account for only 7% of mice undergoing TBI on E13.5 surviving beyond weaning. It is notable that many of the pathological changes in the brain were rescued upon maternal administration of JP4-039 one full day after the 3 Gy TBI when brain damage had already occurred. This included significant reduction in cerebral hemorrhage, apoptosis, CSF accumulation and preservation of more brain tissue.

Our study further points to involvement of mitochondrial dysfunction in the TBI associated brain injury. TBI is known to cause disruption of mitochondrial respiration, with oxidative stress ensuing (*27–30*) with increased ROS production. JP4-039 is a gramicidin S (GS)- derived nitroxide that can convert to the corresponding hydroxylamine after scavenging an electron and a proton from the electron transport chain, thus detoxifying peroxyl and alkoxyl radicals and cycling back to nitroxide to complete a catalytic turnover. JP4-039 could be transported cross placental-fetal barrier to reach fetuses and this transport process was not disturbed by 3Gy irradiation. Using the novel 4D Oxy-wavelet MRI to probe *in vivo* mitochondrial function, we showed 3Gy TBI resulted in mitochondrial respiration dysfunction that was further verified by *ex vivo* Oroboros mitochondrial respirometry. However, upon maternal administration of a single dose of JP4-039 one day after TBI, mitochondrial respiration was rescued concomitant with reductions in fetal brain injury, cerebral hemorrhage, brain tissue loss, and CSF accumulation. The single maternal administration of JP4-039 at mid-gestation also greatly improved long term postnatal survival. Together these findings suggest ROS damage from the perturbation of mitochondrial respiration plays a key role in TBI induced brain damage.

TBI has been shown to cause multi-organ systems injury and prenatal lethality that may include not only mitochondrial dysfunction and oxidative stress, but also other pathogenetic mechanisms. Thus treatments targeting inflammation (*55–57*), cellular redux metabolism (*58*), and thrombosis with the thrombopoietin mimetic JNJ-26366821 (*59*), showed mitigation of TBI injury in adult TBI. Given the important role of mitochondrial in the regulation of cell death pathways, it is worth noting that treatment of JP4-039 also has been shown to reduce apoptosis and also ferroptosis (*60*) after TBI in adult animals. Other drugs targeting necrosis also were shown to further improve outcomes after TBI (*41*). Given the *in vivo* half-life (t_1/2_) of JP4-039 is ∼ 4 hours (*46, 61*), and we observed improved mitochondrial function on E16.5, this would suggest JP4-039 can trigger long lasting changes beyond stochastic scavenging of ROS by JP4-039. We previously showed JP4-039 can stabilize cytochrome c/cardiolipin, consequently reducing phospholipid peroxidation to reprogram the cellular redux state (*62–64*).

Overall, our findings highlight mitochondria as a new therapeutic target for mitigating fetal brain injury from TBI, with JP4-039 as a potential therapeutic agent for *in utero* intervention for mid-gestation TBI.

The *in utero* impact of TBI is difficult to investigate clinically in human pregnancies, making the study of animal models indispensable. This is especially important in the development of therapy, where mechanistic insights are essential for identification of druggable targets. Investigations requiring longitudinal *in vivo* analysis of pregnancies are especially hampered by fetal motion artifacts, making it necessary for most fetal TBI studies to be conducted *ex vivo* in isolated tissue or fixed fetuses. With such ex vivo analyses, fetal-placental interactions and the temporal progression of disease arising from in utero TBI are difficult to study. We note TBI causes injury not only to the heart and brain, but all organs are impacted, though this has not been investigated in the present study. Our novel *in vivo* 4D-uMRI is time- and motion-resolved, and can provide longitudinal monitoring with multi-organ fetal imaging, and the imaging of fetal-placental interactions to track TBI progression, and potential rescue with therapeutic interventions. In addition, *in vivo* 4D-uMRI will allow 4D imaging of all the fetuses in an entire litter, making it possible to interrogate phenotype-genotype correlations that can be used to explore mutations that can impact TBI injury responses and determine susceptibility to fetal TBI. The novel *in vivo* 4D Oxy-wavelet MRI also can provide a unique tool for evaluating *in vivo* mitochondrial function, with the automatic time-and-frequency waveform analysis providing an automated computational pipeline as a surrogate biomarker for high-throughput drug screens in preclinical studies evaluating therapeutic compounds for efficacy in rescuing fetal TBI induced injury across gestational stages.

In summary, we showed the cause of postnatal lethality from in utero TBI is likely from severe brain injury, and with JP4-039 treatment shown to be effective as a radiation mitigator, showing efficacy in rescuing the TBI induced brain damage and subsequent postnatal lethality. These findings confirm and extend our previous studies, further demonstrating the TBI induced lethality involves mitochondrial respiration defect, and with JP4-039 (*65*) treatment showing mitigation of the TBI lethal effects with rescue of mitochondrial respiration. We showed JP4 not only can pass through the placental barrier, but also the blood brain barrier to rescue fetal brain mitochondrial respiration and brain radiation injury. As JP4-039 administration did not change litter sizes or fetus viability, this suggests JP4-039 can be deployed as a safe and effective mitigator of injury from mid-gestational ionizing radiation exposure. These findings suggest JP4 can be an effective mitigator of catastrophic radiation exposure from a terrorist attack in the unborn fetus, with administration even one day after exposure able to affect rescue. Further studies are warranted to investigate the precise window of time when JP4-039 may be administered and affect rescue of conceptuses from in utero radiation exposure. Whether JP4-039 may be given prophylactically before radiation exposure to provide protection from subsequent radiation exposure is another important question worthy of further investigation in future studies.

## MATERIALS AND METHODS

### Mice and Animal Housing

Six-to-eight-week old inbred C57BL/6NTac mice were obtained from Taconic Biosciences b (Model B6). All animals received humane care in compliance with the Guide for the Care and Use of Laboratory Animals (*66*). All experiments were conducted with the approval by the University of Pittsburgh Institutional Animal Care and Use Committee (IACUC). All animals were housed in the AALAS certified animal vivarium maintained on standard laboratory chow, deionized water, and housed 1 breeding pair or litter per cage according to Institutional IACUC protocols (*44*). Mice were provided with food and water ad libitum and kept on a 12:12 hour dark/light cycle. Timed pregnancy was determined by the presence of vaginal plugs after crossing a male and a female in the same cage overnight. Total 76 pregnant females with fetuses were examined, including 24 which received 3 Gy irradiation, 27 which received 3 Gy irradiation plus JP4-039 treatment, 19 mice received 0 Gy irradiation, and 6 mice received 0 Gy plus JP4-039 treatment.

### Total Body Irradiation (TBI)

Pregnant female mice (n=51) were irradiated with 3 Gy irradiation on E13.5 using a Cesium Gamma cell irradiator dose rate 290 cGy/minute according to the published procedures (*67*). Sham controls (n=25) went through the same procedure but with 0 Gy irradiation.

### Administration of Radiation Mitigator GS-Nitroxide, JP4-039

The radiation mitigator JP4-039 (20 mg/kg) was administered at E14.5 to pregnant female mice intravenously in 30% (2-hydroxypropyl)-β-cyclodextrin.). Control groups included pregnant female mice receiving formulation alone or no drug administration. The radiation mitigator was delivered intravenously to pregnant female mice (n=27) on E14.5 at 24 hours after 3 Gy TBI on E13.5.

### Quantitation of JP4-039 in Fetuses, Placentae, and Maternal Plasma with Liquid chromatography–mass spectrometry (LC-MS)

Two groups (N=3 each) of E14.5 pregnant female mice were dosed by intravenous tail vein injection of JP4-039 at 20 mg/kg in 30% (2-hydroxypropyl)-β-cyclodextrin vehicle (2 mg/mL) 24 h after being sham-irradiated or irradiated at 3.0 Gy total body irradiation. Mice were euthanized by CO_2_ inhalation at 30 min after dosing and the following tissues were collected: maternal blood and brain, and for each fetus the placenta, brain, and body. Blood was collected by cardiac puncture using EDTA anticoagulated needles and syringes, transferred to microcentrifuge tubes, and centrifuged at 12,000 x g for 4 min to obtain plasma. Tissues were weighed and all samples were flash frozen using liquid nitrogen and stored at −80 °C until analysis. Samples were analyzed with a validated LC-MS/MS assay as published (*72*).

### Analysis of Live Births, Newborn Growth and Development

Twenty-eight pregnant female mice were followed for time of delivery of litters, litter size, the number of mice per litter, survivors to birth, as well as a growth of surviving mice to weaning, including 10 mice that received 3 Gy irradiation, 10 mice that received 3 Gy irradiation plus JP4-039 treatment, 5 mice that received 0 Gy irradiation, and 3 mice that received 0 Gy plus JP4-039 treatment.

### *In vivo* Ultrasound and Doppler Echocardiography

*In vivo* ultrasound was performed on E15.5 with our routine protocols (*68–70*) using a Fiji Film/Visual Sonic Vevo 2100 scanner with a 40 MHz transducer. The imaging modalities used included B-mode imaging, color flow and spectral Doppler, and M-mode imaging as previously described to evaluate fetal structural heart defects. For left ventricular (LV) functions, the areas of the LV chamber in diastole (LVEDA) and systole (LVESA) were measured, and the fraction area change (FAC%) was calculated as (LVEDA – LVESA) × 100/LVEDA. The foramen ovale was imaged by color-flow Doppler in four-chamber view.

### Free-breathing no-gating 4D *in utero* MRI (4D-uMRI)

This novel time-and-motion resolved 4D *in utero* MRI (4D-uMRI) is capable of simultaneous *in utero* anatomical and functional imaging of multiple mouse fetuses in a single all-encompassing scan with both high spatial [0.0017 mm^3^] and temporal (∼14 msec) temporal resolutions without motion artifact. It is enabled by a low-rank and sparse modeling of a 4D image ρ(**r**, *t*) (for 3D spatial position **r** and time *t*). The low-rank model expresses the image as the product of *L* spatial coefficient maps 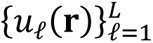 and temporal basis functions 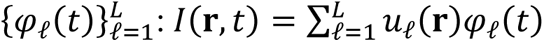. This approach exploits the correlation of images over time, which also can be combined with a transform-sparsity model of the image series to allow imaging with high spatiotemporal resolution. Memory-efficient image reconstruction is possible by reframing image reconstruction as direct recovery of model parameters (i.e., the spatial coefficient maps and temporal basis functions). Whereas conventional fast MRI suffers from motion artifacts, this novel 4D-uMRI (fast time series of isotropic 3D stacks) can model fast respiratory, cardiac, and fetal motion, thus enabling free-breathing-no-gating motion-resolved 4D-uMRI for non-invasive fetal imaging to track in utero development of the mouse brain throughout mid- to late-gestation.

### *In vivo* 4D-uMRI Acquisition

Free-breathing gating-free *in vivo* 4D-uMRI was performed on E16.5 in a blinded manner. MRI operators and analyzers were blinded to the identity of the mice. All pregnant female mice (n= 5 per group) received general inhalation anesthesia with Isoflurane for *in vivo* MRI. The mice were placed into a clear plexiglass anesthesia induction box that allows unimpeded visual monitoring of the animals. Induction was achieved by administration of 3% Isoflurane mixed with oxygen for a few minutes. Depth of anesthesia was monitored by toe reflex (extension of limbs, spine positioning) and respiration rate. Once the plain of anesthesia was established, it was maintained with 1-2 % Isoflurane in oxygen via a designated nose cone and the mouse was transferred to the designated animal bed for imaging. Respiration was monitored using a pneumatic sensor placed between the animal bed and the mouse’s abdomen while rectal temperature was measured with a fiber optic sensor and maintained with feedback-controlled warm air source (SA Instruments, Stony Brook, NY, USA). Oscillating acute hypoxia challenges during 4D u-MRI acquisition were applied via the nose cone by intermittent titrating breathing air with 10% oxygen with 90% nitrogen or 100% oxygen every 3 minutes.

*In vivo* 4D-uMRI was carried out on a Bruker BioSpec 70/30 USR spectrometer (Bruker BioSpin MRI, Billerica, MA, USA) operating at 7-Tesla field strength, equipped with an actively shielded gradient system and a quadrature RF coil with an inner-diameter of 35 mm with the following parameters: field of view (FOV) = 4.0 cm X 2.78 cm X 2.18 cm, acquisition matrix = 256 × 178 × 140, isotropic resolution = 156 μm X 156 μm X 156 μm, echo time (TE) = 2.98 msec, repetition time (TR) = 6.687 msec, flip angle (FA) = 10^0^.

Fetal brains and placentas were manually segmented by blinded analyzers on the reconstructed 4D-uMRI using the open source ITK-Snap (http://www.itksnap.org). The regions of interest (ROIs) were used for volumetric and dynamic signal analysis.

### *In vivo* 4D Oxy-wavelet MRI for Probing Mitochondrial Functions

Our 4D-uMRI is not only time- and motion-resolved, but functional signals can also be captured in the same 4D scan. This allows quantifying fast blood-oxygen-level-dependent (BOLD)(*71–74*) dynamics, the basis for functional MRI (fMRI)(*75–82*). In conjunction with oscillating short bursts (3 min) of hypoxia (10% O_2_) challenges (Fig.6A), we have successfully established 4D Oxy-wavelet MRI (4D-uMRI with oscillating hypoxia challenges) to probe mitochondrial function *in vivo,* leveraging the fact that acute response to hypoxia challenges require the transient, reversible, compensatory activation of the mitochondrial respiratory chain (*47–53*) to maintain oxygen homeostasis.

To aid in automating the diagnosis, we developed an automated time-frequency waveform analysis scheme for 4D Oxy-wavelet MRI (Fig.6 L-O). Our BOLD response signals (Fig.6L) are event-driven, following the oscillations of input oxygen cycling; the event oxygen waveform resembles a square wave function (Fig.6A). Cross-correlating the BOLD response signals (Fig.6L) by template event waveforms at several oscillation periods, tau, (Fig.6M) will produce a feature space revealing various details (Fig.6N). The time-Oxy-wavelet space (Fig.6N) is a two-dimensional matrix that contains the value of the correlation at that time point, for that oxygen period. It is comparable to a spectrogram or other time-frequency analysis. The y-axis of that space is time whereas the x-axis is the Oxy-wavelet period, tau. The final Oxy-wavelet index is calculated as the maximum absolute value of the cross-correlation, where the absolute maximum of both marginals is achieved. The location of the peak-magnitude value reveals the midpoint of the oxygen cycling experiment and the oscillation period (neither of which is precisely known *a priori* when oxygen switching is manually triggered). More importantly, the sign of the peak value reveals the direction of oxygen response: a positive peak indicates a positive correlation with the oxygen cycling (tissue hypoxia when external oxygen is decreased), the passive inability to cope with the acute cyclic hypoxia; whereas a negative peak indicates an opposite behavior (brain sparing wherein tissue over-corrects for external oxygen decreases by adjusting to an even higher oxygenation), the ability for active compensation of the acute cyclic hypoxia. Therefore, fetal brains with good mitochondrial respiratory functions exhibit negative Oxy-wavelet indexes, whereas positive Oxy-wavelet indexes indicate poor mitochondrial respiratory functions.

### Ex vivo MRI

Immediately following *in vivo* 4D-uMRI, pregnant females were euthanized to harvest fetuses. One fetus per litter was used for Oroboros mitochondrial respirometry, whereas the rest of the features were fixed with 10% formalin for high-resolution *ex vivo* MRI. Well-fixed mouse fetuses were immersed in 1:200 MultiHance (gadobenate dimeglumine, 529 mg/ml, Bracco Diagnostic, Inc. Monroe Twp, NJ) diluted with phosphate-buffered saline (PBS) at 4^0^C for 48 hours. After staining, mouse fetuses were secured on a tongue depressor (McKesson Medical-Surgical, Irving, TX) with Webglue surgical adhesive (n-butyl cyanoacrylate, Patterson Veterinary, Devens, MA). The fetuses were then immersed in Fomblin Y (perfluoropolyether, Sigma-Aldrich Millipore) to eliminate the susceptibility artifact at the tissue-air interface and to avoid dehydration during imaging. MRI was carried out on a Bruker Biospec 7T/30 system (Bruker Biospin MRI, Billerica,MA) with a 35-mm quadrature coil. 3D MRI was acquired with a fast spin echo sequence, the Rapid Acquisition with Refocusing Echoes (RARE), with the following parameters: field of view (FOV) = 3.99 cm X 1.84 cm X 1.17 cm, acquisition matrix = 1024 × 400 × 140, isotropic resolution = 39 μm X 46 μm X 46 μm, effective echo time (TE) = 24.69 msec, RARE factor 8, repetition time (TR) = 900 msec, and flip angle (FA) = 180^0^. Following 3D imaging reconstruction, the open-source Horos (https://horosproject.org/) software was used for re-orientation of each fetus for visualization.

### Oroboros mitochondrial respirometry on fetal brain lysates

A fast 2D gradient echo MRI with self-gating was performed at the same imaging session on E16.5 to quickly parcellate fetuses into high and low functional states with the following parameters: slice thickness 0.7 mm, field of view (FOV) = 4.0 cm X 4.0 cm, acquisition matrix = 256 × 256, in-plane resolution = 156 μm X 156 μm, echo time (TE) = 1.872 msec, repetition time (TR) = 162.75 msec, and flip angle (FA) = 180^0^, scan time = 5 min 14 sec. Immediately following *in vivo* MRI on E16.5, fetal brains were harvested according to their position on MRI images. One fetus per 3 Gy and 0 Gy groups, and 2 fetuses per 3 Gy with JP4-039 group with one high and one low function were harvested for Oroboros mitochondrial respirometry.

Freshly harvested fetal mouse brains were homogenized in ice-cold Mir05 respiration buffer using a dounce homogenizer. The Mir05 buffer contained 110 mM sucrose,0.5 mM EGTA, 3 mM MgCl_2_, 60 mM K-lactobionate, 20 mM taurine, 10mM KH_2_PO_4_, 20 mM HEPES, and 1g/L fatty acid free BSA. After centrifugation at 600xg to remove debris, the supernatant containing ∼150-200 µg of protein was added to Mir05 respiration medium maintained at 37°C in sealed Oroboros Oxygraph 2K chambers (Oroboros Instruments, Inc. Innsbruck, Austria). Then, substrates and inhibitors were added in a stepwise fashion as follows. First, cytochrome-C (10 µM) was added to limit artifacts caused by damage to the outer mitochondrial membrane during homogenization. Then, malate (2 mM) was added as a TCA cycle substrate followed by ADP (5 mM) to initiate oxidative phosphorylation (State 3 respiration). Pyruvate (5 mM) and glutamate (10 mM) were added to stimulate Complex I, followed by succinate (10 mM) to drive respiration through Complex II. Maximal, uncoupled respiration was initiated with addition of CCCP (1 µM). Finally, rotenone (0.5 µM) was added to inhibit complex I, thereby providing an assessment of the relative contributions of Complex I and Complex II. All respiration data were normalized to total protein concentration.

#### Histopathologic Evaluation

Newborn pups were sacrificed by decapitation using a pair of surgical scissors. Heads were drop-fixed in Bouins’s solution for 48 hours then transferred to 10% neutral buffered formalin for 48 additional hours, sectioned sagittally, placed in cassettes, dehydrated in a graded series of alcohol, and embedded in paraffin blocks. Paraffin-embedded sections were cut to 5µm using a microtome, mounted on glass slides, and stained with routine hematoxylin and eosin (H&E).

#### Fluorescent Microscopy for Cell Death

Three fetal brains per group were harvested for cell death evaluation with fluorescent microscopy. Apoptosis staining was done using TMR red In Situ Cell Death Detection Kit (121567910, Roche). Brain sections (5 um) were incubated with reaction mixture at 37 °C for 45 minutes followed by 3 washes of PBS. Sections were incubated with 1:500 dilution Alexa 488 Flash Phalloidin (424201, Biolegend) for 45 minutes at room temperature. Sections were washed 3 times with PBS, incubated one minute in Hoechst dye (B2883, Sigma) at 1 mg/100mL in dH_2_O, followed by 3 PBS washes and coverslipped with gelvatol.

#### Statistical Evaluation

All data were summarized with mean ± standard deviation (SD). For data on maternal mice, values were compared between groups with the two-sided t-test.

## AKNOWLEDGEMENT

This study was supported by NIAID/NIH grant U19-A168021, in part by award R50-CA211241, and in part by funding to YLW from NIH (EB023507, NS121706-01), AHA (18CDA34140024), and DoD (W81XWH1810070, W81XWH-22-1-0221). This project used the UPCI animal facility and the Cancer Pharmacokinetic and Pharmacodynamic Facility (CPPF) which are supported in part by award P30CA047904. This project also used the Rangos Research Center Animal Imaging Core of the Children’s Hospital of Pittsburgh of UPMC for MRI.

## Supplemental Materials

**Supplemental Figure. Fetal cardiac function on E15.5 evaluated by *in vivo* fetal ultrasound. 3Gy TBI on E13.5 did not cause significant changes in fetal cardiac function.**

(A) left ventricular ejection fraction (LV EF %) and left ventricular fractional shortening (LV FS %). (B) fetal heart stroke volume (SV, ul). (C) fetal cardiac output (CO, ul/min). (D) fetal heart rate (HR, bpm). Red-naïve control, Green – 3 Gy, Blue – 3Gy + JP4-039.

