## Supplementary figures and images for "Mitigation of Fetal Irradiation Injury from Mid-Gestation Total Body Radiation with Mitochondrial-Targeted GS-Nitroxide JP4-039"

### Supplemental Figure 1

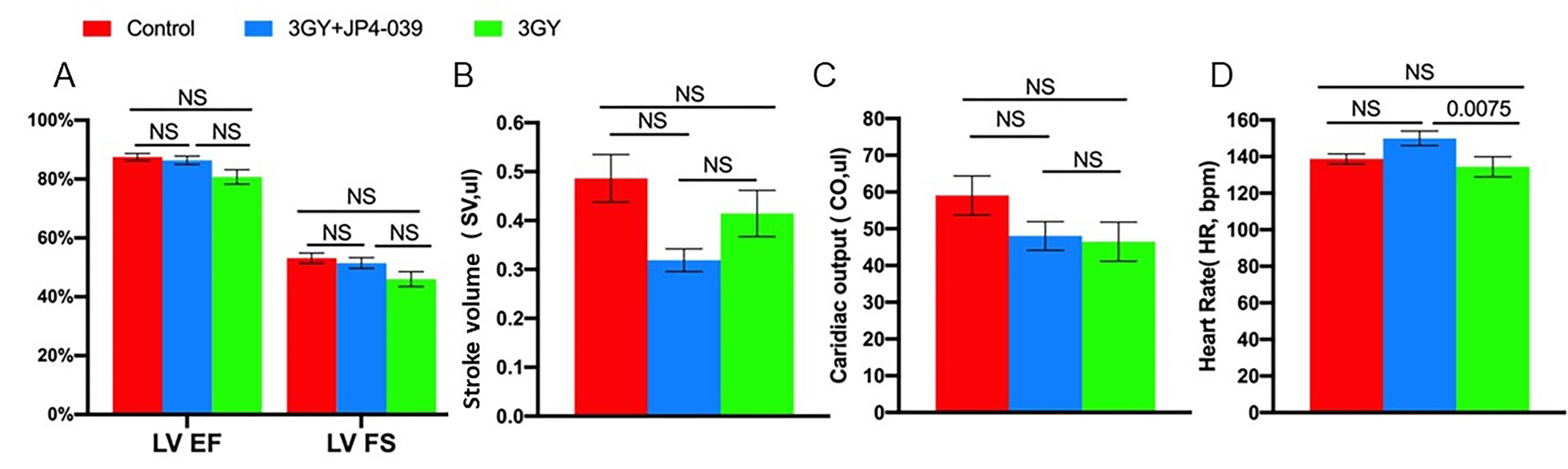
